# Mapping Tissue-Specific Protease Dynamics in the Pig Respiratory Tract During Influenza A Virus Infection

**DOI:** 10.64898/2026.02.11.705325

**Authors:** Chrysillis Hellemann Polhaus, Aleksander Moldt Haack, Ramona Trebbien, Lars Erik Larsen, Ulrich auf dem Keller, Kerstin Skovgaard, Konstantinos Kalogeropoulos

## Abstract

Influenza A virus (IAV) is a zoonotic pathogen capable of infecting diverse avian and mammalian hosts, causing seasonal epidemics and occasional global pandemics in humans. Viral entry requires proteolytic activation of hemagglutinin (HA). While serine proteases such as TMPRSS2, and HAT are known HA activators, the respiratory tract harbours additional proteases whose contributions to infection remain unclear. Dysregulation of these proteases can enhance viral replication, tissue damage, and inflammation, highlighting the need for a systems-level view of the proteolytic landscape. Here, we use high-throughput, proteome-wide proteomics and N-terminomics to identify 112 host proteases across the nasal mucosa, trachea, and lung. We monitor and validate 28 proteases with targeted proteomics and microfluidic qPCR, representing a comprehensive degradome analysis in the respiratory tract of the highly translational pig model of influenza infection. We show that protease abundance and activity were highly tissue-specific: while the nasal mucosa showed selective activation of broad- and narrow-specificity proteases alongside robust antiviral responses, the trachea exhibited modest modulation with subtle shifts in protease–inhibitor balance, and the lung maintained predominantly active proteases despite lower viral loads but severe tissue damage, indicative of immune-mediated pathology. Sequence motif analysis revealed distinct cleavage preferences across tissues, indicating differential protease processing across the studied respiratory tissues in antiviral pathways, antigen processing, and tissue remodelling. Several identified proteases, including ST14, KLKB1, PRSS8, and LGMN, were increased and functionally active upon infection, suggesting roles in viral processing and host immune regulation. Collectively, our results define a spatially organised proteolytic network that shapes tissue-specific antiviral host responses and contributes to H1N1 influenza pathogenesis.

## Introduction

Influenza A virus (IAV) is a zoonotic virus that can infect a wide range of animals, including avian and mammalian species. IAV causes respiratory illness worldwide in humans, leading to annual seasonal epidemics and occasional global pandemics^1^. All IAV pandemics have been caused by zoonotic transmission from avian or swine sources, followed by reassortment and emergence of novel IAV strains after adaptation to a new host. IAV is an enveloped negative-sense RNA virus that contains eight segments that encode at least 10 proteins. Segment four encodes the viral surface protein hemagglutinin (HA) that facilitates viral entry by binding to sialic acid receptors on host cells and mediating membrane fusion during endocytosis^1^. HA is produced as an inactive precursor, HA0, which must be cleaved into HA1 and HA2 by host proteases^2^. This activation triggers a conformational change that enables viral envelope fusion with the endosomal membrane, allowing viral ribonucleoprotein (vRNP) release into the host cytoplasm. Subsequently, the vRNP is transported into the host nucleus, where viral replication and transcription are initiated^2^.

Three serine proteases, TMPRSS11D (HAT), TMPRSS2, and KLKB1 are known to facilitate the cleavage of HA^3–9^. Serine proteases function as endopeptidases by cleaving peptide bonds within the protein. Their active site contains a serine, which is part of a catalytic triad along with histidine and aspartate, enabling the hydrolysis of peptide bonds. The preferred cleavage sites vary among different serine proteases. While HAT and TMPRSS2 are well-established as critical host proteases for HA activation in influenza virus infection, the complex proteolytic landscape in the respiratory tract, along with host-specific responses to different IAV strains provides a strong rationale for exploring whether additional proteases are directly or indirectly involved in HA activation or host responses to IAV infection.

Proteases in the respiratory tract are crucial in maintaining tissue homeostasis, regulating immune responses, and supporting normal respiratory function. Under healthy conditions, protease activity is tightly controlled to prevent excessive tissue damage and maintain epithelial integrity. However, during influenza infection, dysregulation of the protease network can facilitate viral entry, enhance viral replication, and contribute to pathological changes such as inflammation, epithelial damage, and impaired lung function^10^. The serine proteases KLKB1, F2, and PLG are among the most common proteases found in the lung^4,11,12^, while the serine protease TMPRSS2 is commonly found in the proximal airways^7,8^. Other host proteases, such as cathepsins and matrix metalloproteinases, have also been found previously in respiratory tissues^10^.

A deeper understanding of how host proteases shape viral infection and influence multiple host pathways requires a comprehensive, systems-level perspective. Proteome-wide profiling of protease activity through N-terminomics enables mapping of the respiratory tract’s proteolytic landscape in both healthy and diseased states. By capturing newly generated protein N-termini produced by proteolytic cleavage, this approach provides a global view of protease activity rather than protein abundance alone. Mass-spectrometry-based analysis of these cleavage events reveals whether proteolysis activates functional proteins or represent dysregulated processing, which is a hallmark of disease.

In this study, we mapped the host protease landscape during IAV infection using N-terminomics, providing a system-wide view of protein and protease network dynamics in response to a pre-pandemic H1N1 strain. We identified 112 proteases across the nasal mucosa, trachea, and lung of pigs three days post-infection and validated these findings using microfluidic qPCR and targeted proteomics (PRM), representing the first such analysis in pig respiratory tissues. H1N1 elicited a classical antiviral response in the upper airways with concurrent tissue-specific protease activity. Tightly regulated in the nasal mucosa, modest in the trachea, and predominantly activity-driven in the lung. Overall, our results map tissue-specific protease dynamics during influenza virus infection, providing a foundation to understand how the host protease network orchestrates antiviral defence and contributes to respiratory pathology.

## Results

### Influenza A virus induces a classical innate immune response in the porcine model

IAV infection triggers complex host proteolytic and immune responses that vary across respiratory tissues. To capture these dynamics, we profiled the protease landscape and full proteome in distinct respiratory compartments of pigs following IAV infection. We included tissue from the nasal mucosa, the mucosal membrane from the trachea, and lower lung tissue (*lobus dexter caudalis*) was used for HUNTER and global proteome analysis. The experimental design is shown in Fig. 1A. After infection, we measured the quantity of viral RNA (Fig. 1B), where the highest viral load was seen in the nasal mucosa tissue (average 1.41 ∗ 10^6^ copies/mL), the trachea (average 1.06 ∗ 106 copies/mL), and the lowest viral load was seen in the lung (average 6.25 ∗ 10^3^ copies/mL).

**Figure 1.**
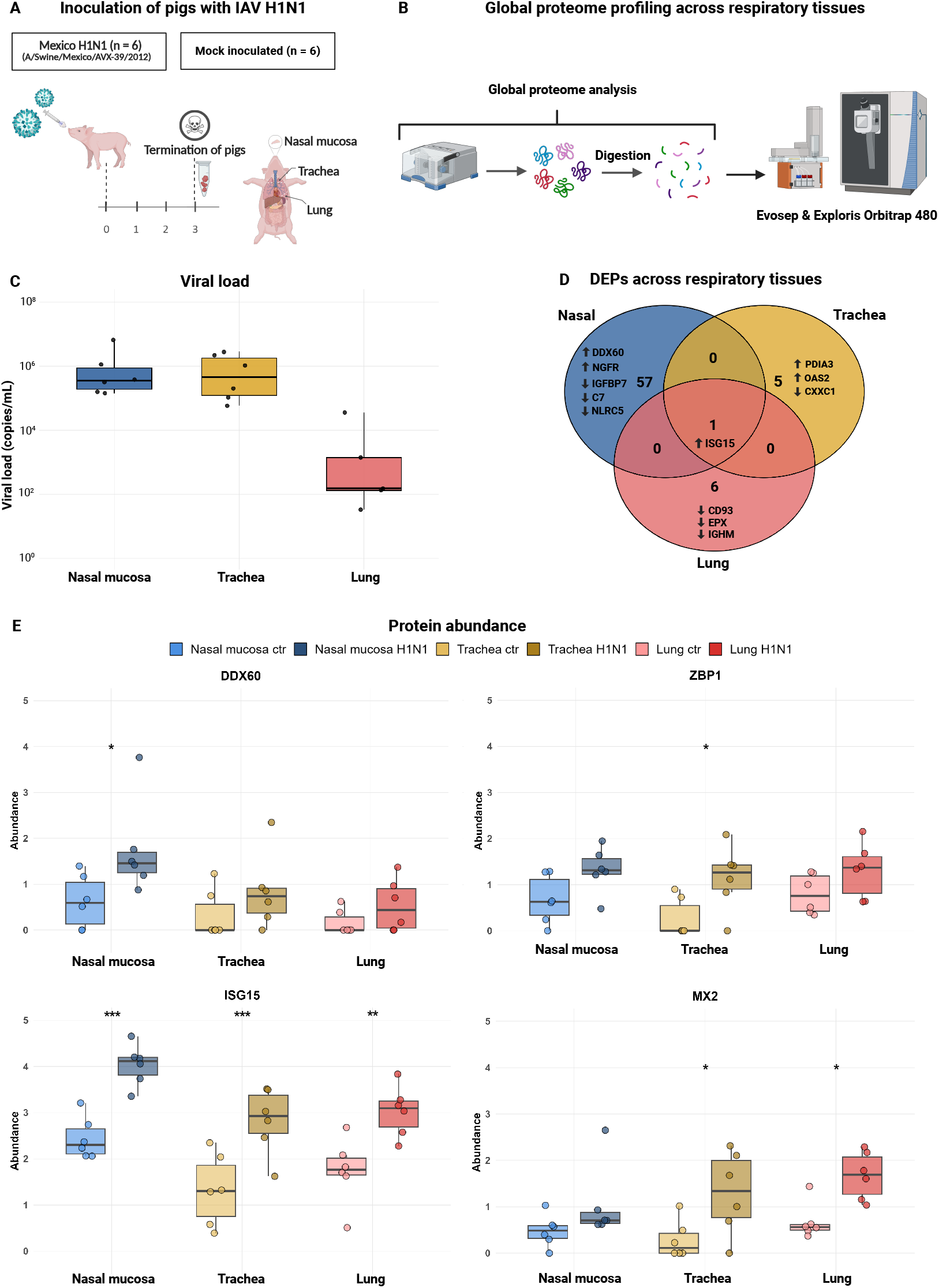
Global proteome analysis revealed an antiviral innate immune response in the respiratory tissue post IAV H1N1 infection. A. Experimental design. 6 pigs were inoculated with IAV H1N1, and 6 pigs were mock inoculated. They were euthanized 3 days post infection (dpi). Three different respiratory tissues were used for analysis, created in Biorender. B. Experimental design for global proteome analysis, samples were analysed on Evosep and Exploris 480 for 58 min, created in Biorender. C. Viral load, RNA viral load in nasal mucosa, trachea, and lung tissue 3 DPI. Pig 32 had no viral load in the lung and is not shown in the figure. D. Venn diagram of the total number of DEPs in each tissue. E. Boxplots of 4 different antiviral proteins, DDX60, ZBP1, ISG15 and MX2. SEM is depicted by error bars. Statistical significance: p < 0.1 (*), p < 0.05 (**), p < 0.01 (***).

We identified a total of 5,571 protein groups and 54,352 peptides in our control and infected proteome samples. Of these, 4,805 protein groups were consistently quantified across all three tissues included in the study (Supplementary Table S4). The highest number of differentially expressed proteins (DEPs) was seen in the nasal mucosa tissue of control vs infected pigs, where 57 DEPs were found. The well-characterised antiviral interferon-stimulated genes (ISGs), ISG15 and OAS2, were among the proteins that were elevated in one or several respiratory tissues of the infected pigs.

We performed enrichment analysis of the DEPs in the nasal mucosa, trachea, and lung tissues by comparing control and H1N1-infected pigs. Two types of analyses were conducted: gseKEGG and Gene Ontology (GO) (Supplementary Table S5 and Supplementary Table S6). In the nasal mucosa, our enrichment analysis identified activation of the Influenza A pathway, endocytosis and NOD-like receptor signalling pathway, in H1N1-infected tissue. We identified a similar set of pathways in the trachea, albeit their regulation was less pronounced compared to the nasal mucosa. Notably, Influenza A and coronavirus disease – COVID-19 were among the enriched pathways. Enrichment analysis of ontology in the trachea also indicated strong associations with infection, as the pathway: response to stress, which was one of the pathways activated after infection in the trachea. For the lung tissue, our analysis found enriched pathways related to infection, including Toll-like receptor signalling and PI3K-Akt signalling pathway.

Several key ISGs, including ISG15 and OAS2, showed increased abundance following IAV infection (Fig. 1E). ISG15 is a ubiquitin-like protein that plays a critical role in antiviral defence by modifying host and viral proteins to limit viral replication^13,14^. At the same time, OAS2 activates RNase L to degrade viral RNA, making both essential effectors/indicators of the host’s innate immune response^13^. DDX60, which promotes RIG-I-like receptor signalling^15^, was elevated specifically in the nasal mucosa. MX2 and the viral pattern recognition receptor ZBP1 were more abundant in the trachea, suggesting a localised antiviral response in which MX2 may contribute to restriction of viral replication through interference with viral nucleocapsid trafficking. In contrast, several ribosomal proteins involved in viral mRNA translation and other immune regulators were decreased, particularly in the nasal mucosa and lung, highlighting tissue-specific antiviral and immune responses across the porcine respiratory tract.

Comparison of control and infected pigs revealed tissue-specific remodelling of the extracellular matrix (ECM). In the nasal mucosa, several ECM glycoproteins and proteoglycans (e.g., COL5A2, BCAN) were increased, indicating active tissue reorganisation, while integrins and adhesion proteins (e.g., ITGAV, THBS2) were decreased, suggesting loosening of cell–matrix contacts that may facilitate immune cell infiltration (Supplementary Table S4). In the trachea, increased levels of folding and glycosylation enzymes (e.g., PDIA3, GALNT4) coincided with decreased structural cytoskeletal components (e.g., COL1A1, KRT2), consistent with ECM turnover and epithelial remodelling. In the lung, ECM-associated and adhesion proteins (e.g., RACK1, CD93) were predominantly decreased following infection, reflecting widespread matrix disruption and potentially compromised tissue integrity (Supplementary Table S4). IAV infection triggers tissue-specific host responses across the porcine respiratory tract, with the nasal mucosa exhibiting the most pronounced antiviral and immune activation. Interferon-stimulated proteins and ECM regulators were differentially modulated among tissues, revealing the spatial complexity of host defence mechanisms. Together, these findings provide a comprehensive characterisation of the early immune landscape in IAV-infected pigs, demonstrating a rapid induction of innate immune pathways and matrix remodelling processes.

### N-terminomics analysis reveals a tissue-specific protease landscape in the respiratory tract

Our global proteome analysis identified 112 proteases (Supplementary Table S7), suggesting a widespread involvement of proteolytic activity during the host immune response to H1N1 IAV infection. To investigate the protease activity during infection, we analysed N-terminal enriched samples (PO samples) to quantify the substrate activity across all three respiratory tissues (Fig. 2A).

**Figure 2.**
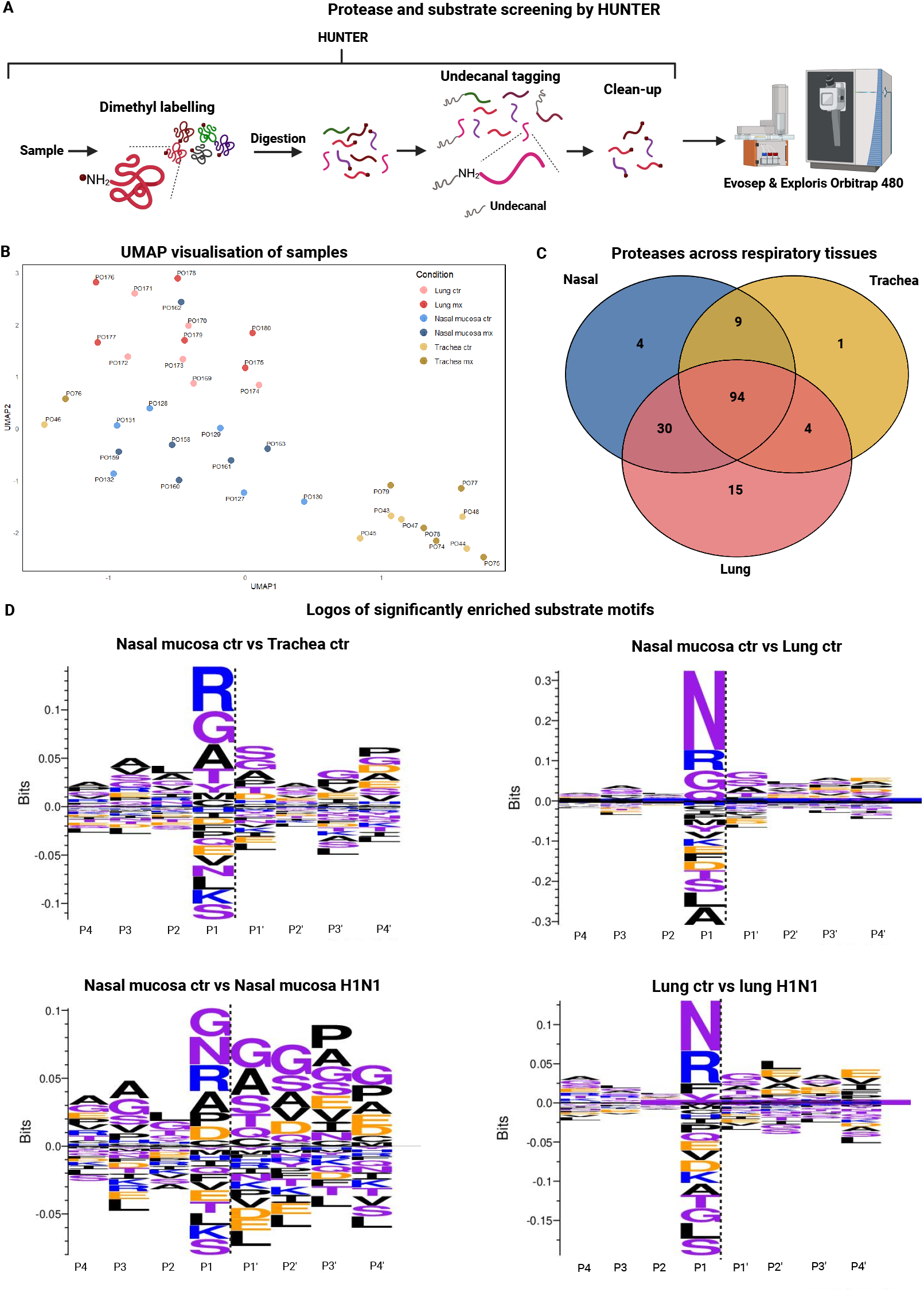
N-terminomics analysis reveals significant cleavage patterns in lower respiratory tissues. A. Experimental design. Three different respiratory tissues were used for HUNTER. Samples were analysed on Evosep and Exploris 480 for 58 min, created in Biorender. B. UMAP of enriched samples; blue colours indicate nasal mucosa tissue, yellow colours indicate trachea tissues, and purple colours indicate lung samples. C. Venn diagram of the union of proteases quantified in the three tissues. D. Sequence logos of cleavage observed in the three tissues, control vs control, control vs H1N1 infected tissue. The logos presented here are based on significantly regulated peptides (adjusted p < 0.1). The remaining logos of Trachea ctr vs Lung ctr is available in Supplementary Figure S1. Sequence logos were made using Seq2Logo.

UMAP analysis of all samples (Fig. 2B) revealed clear clustering of proteolytic activity by tissue type, while separation by infection status was less pronounced, indicating that tissue-specific expression patterns primarily define the protease substrate landscape. Most samples clustered by tissue, although two tracheal samples (one control, PO46; one infected, PO76) grouped with the nasal mucosa, indicating minor inter-tissue variability. We initially characterised the overall protease landscape in the three tissues, identifying 112 proteases. However, not all proteases were quantified in every tissue. Of the 112 proteases, 53 are exopeptidases, 58 are endopeptidases, and one exhibits dual functionality (Supplementary Table S7). Exopeptidases are distributed across multiple protease classes, whereas endopeptidases are predominantly serine and cysteine proteases, with relatively few metalloproteases. While most proteases were detected in the lung, a large proportion were shared across the nasal mucosa and trachea, highlighting a dynamic yet broadly overlapping tissue-specific protease landscape (Fig. 2C). Twenty-eight proteases were shared between the nasal mucosa and lung, while twelve proteases were uniquely detected in the lung tissue. Interestingly, DPP7 was the only protease detected in both the nasal mucosa and trachea, but it was absent in the lung.

### Tissue-specific protease remodelling during H1N1

To investigate tissue-specific protease regulation in the porcine respiratory tract, we first analysed healthy control samples and H1N1-infected tissues. We combined global proteome and N-terminomics approaches to capture both total protein abundance and protease activation states. This strategy allowed us to assess steady-state differences across the nasal mucosa, trachea, and lung, and to reveal localised, tissue-specific changes in protease activity and inhibitor abundance in response to viral infection. To examine tissue-specific protease expression under steady-state conditions, we analysed healthy control samples and found that the lung contained the largest number of uniquely differentially abundant protease peptides compared to the nasal mucosa and trachea. In the global proteome samples, CFB, CTSB, ELANE, PRTN3, and TGM2 were higher in the lung compared to the trachea. CNDP2, CTSH, SCPEP1, TGM2, and UCHL1 were also more abundant in the lung than in the nasal mucosa.

We also observed the full peptide of UCHL1 being significantly decreased in the lung compared to the nasal mucosa. CTSH cleavage past the second propeptide suggested increased active CTSH in healthy lung tissue relative to nasal mucosa. Whereas several peptides were found significantly increased for TGM2 all which were cleaved at different positions all indicating degradation of TGM2 in the healthy lung tissue. Signal peptide removal was observed for CPB2 and KLKB1 in the healthy nasal mucosa along with a significantly increase of the active TPP1 in the nasal mucosa compared to the trachea. In the analysis of the enriched peptides, most inhibitors were more abundant in the lung than in the nasal mucosa or trachea (A2M, CAST, ITIH4, LXN, PEBP1). PI15 differed only between lung and nasal mucosa, while SERPINA1, ITIH1, and ITIH3 were higher in lung versus trachea. In nasal mucosa versus trachea, ITIH1, ITIH3, and SERPINA1 were elevated, with PI15 higher in the trachea. To further investigate protease recognition motifs and their potential biological significance, sequence logos were generated using Seq2Logo. These logos highlight enriched and depleted amino acids surrounding the cleavage sites. No distinct cleavage motif is apparent in the upper respiratory system, although residues such as arginine (R), glycine (G), and alanine (A) appear moderately enriched (Fig. 2D). In contrast, we find a clear preference for asparagine (N) at the P1 positions in the lung, suggesting a potential substrate specificity for LGMN in these comparisons (Fig. 2D). Overall, our analysis of the healthy tissues reveals tissue-specific patterns of protease and protease inhibitor activity under steady-state conditions. Protease and inhibitor abundance are highly tissue-specific, with the lung showing the greatest diversity. Cleavage site analysis indicates moderate enrichment of arginine (R), glycine (G), and alanine (A), and a clear preference for asparagine (N), suggesting potential substrate specificity.

Following H1N1 infection, we observed tissue-specific changes in protease regulation across all three respiratory tissues. These changes reflect a localised protease activity, processing and inhibitor abundance with implications for cell death, extracellular matrix remodelling and inflammatory responses. Quantitative global proteome analysis revealed a significant decrease in F12 and MMP2 protein abundance in the H1N1-infected nasal mucosa compared to healthy controls (Supplementary Figure S2). We did not observe significant changes in protease abundance in the infected trachea or lung relative to their respective controls.

In the enriched N-terminomics analysis, we observed several protease-derived peptides that displayed altered abundance upon infection, suggesting changes in protease processing or activation states. Specifically, seven protease-derived N-terminal peptides were differentially abundant in infected nasal mucosa compared to control samples, while twenty peptides corresponding to proteases were differentially abundant in the infected lung tissue. In contrast, no protease-derived peptides met the thresholds for differential abundance in tracheal tissue.

In the nasal mucosa, viral infection reduced N-terminal peptides from THOP1 and mature F2, while increasing internal peptides from IDE, trypsin, and SCPEP1. Several protease inhibitors were elevated (SERPINB6, SERPIND1, PI15, PI16, LXN, ITIH4, and others), whereas TIMP3 was reduced.

Twenty protease-derived N-terminal peptides were differentially regulated in lung tissue following H1N1 infection, with trypsin being the only protease decreased post-infection. In contrast, multiple protease inhibitors were increased (A2M, ITIH4, LXN, PI15, PI16, SERPINA3-2, SERPINB6, SERPINB9, SERPINC1, SERPING1, and TIMP2), while only GSK3A and TIMP1 were reduced.

The internal peptide for the protease IDE had a similar pattern across both lung and nasal mucosa, it was seen increased in both tissues after infection. The total protein abundance of IDE was increased in the H1N1 infected nasal mucosa and trachea compared to their respective controls (Supplementary Figures S2). The mature peptide of F2 was seen decreased in the H1N1 infected nasal mucosa, but the opposite was observed in the H1N1 infected lung (Supplementary Figures S2).

Even though no differentially abundant proteases were found in the trachea tissue, some of the proteases were identified (including CASP7, PIGK, and PRSS8). Three different protease inhibitors were also found decreased in the H1N1 infected trachea tissue compared to the controls (e.g. A2M, LXN and PI15).

To understand the cleavage pattern post-infection in each of the tissues, we generated sequence logos for the significant substrates when comparing control vs H1N1-infected in both nasal mucosa tissue and lung tissue (Fig. 2D). We found no clear signature when analysing the nasal mucosa tissue, whereas asparagine (N) and arginine (R) are more predominant in the p1 position in the lung tissue.

Comparing the infected tissues with each other in the global proteome analysis, PRTN3 and TGM2 were significantly differentially abundant in the infected lung tissue compared to the infected nasal mucosa and the infected trachea. We also observed that TGM2 was significantly differentially abundant in the infected trachea compared to the infected nasal mucosa. TMPRSS11D and ANPEP were both significantly increased in the infected nasal mucosa tissue compared to the infected lung. MMP14 and KLKB1 were also significantly increased in the infected nasal mucosa tissue compared to the infected trachea. Although total protease abundance was largely similar across tissues, we found that N-terminomics revealed tissue-specific differences in protease-derived peptides, reflecting localized activation or processing. These results reveal tissue-specific protease activation during infection, with lung, nasal mucosa, and trachea, each exhibiting distinct patterns of protease-derived peptides and reflecting localised processing events (Supplementary Table S8).

Overall, we demonstrate that H1N1 infection primarily modulates protease activation rather than expression. While total protein levels of most proteases were largely unchanged, the N-terminomics analysis revealed localised activation or processing events, with distinct patterns in nasal mucosa and lung. Notably, IDE was consistently activated across nasal mucosa and lung, whereas other proteases, including F12, MMP2, and F2, exhibited tissue-specific regulation. Concurrent changes in endogenous protease inhibitors (SERPINB6, SERPIND1, PI15, PI16, LXN, and ITIH4), suggest coordinated modulation of serine and metalloprotease activity. These proteolytic changes are consistent with localised apoptotic signalling, extracellular matrix remodelling, and inflammatory regulation, highlighting a complex and tissue-specific host response to viral infection.

### Tissue-specific protease activity shapes immune and ECM remodelling during influenza A infection

We used three databases (MEROPS^16^, TopFind^17^, and UniProt^18^) to investigate the identified proteases and their respective cleavages and substrates (Supplementary Table S8). Mapping the/Enrichment analysis of the subsequent data revealed that both nasal and lung tissues share core immune and cellular pathways, including antiviral immunity, translation, and DNA repair. However, the lung also activated pathways related to ECM organisation and viral-host interactions, reflecting its more complex tissue functions.

In the nasal mucosa, cleavage at the P1 position was highly diverse, with glycine (G), asparagine (N), and arginine (R) residues observed (Fig. 2D). Candidate contributors include CAPN1 and CAPN2; ELANE, which preferentially cleaves G; LGMN, favouring N; and CTSB, which prefers R at P1 (Supplementary Table S7). In contrast, lung tissue displayed a more defined profile with a clear preference for N and R. LGMN likely mediates cleavage after N, whereas CTSB and KLKB1 are among the proteases that are predicted to cleave R (Fig. 2D, Supplementary Table S7).

Mapping to Reactome pathways revealed that infection induced localised proteolytic events targeting proteins involved in immune signalling, antiviral defence, apoptosis, stress responses, protein synthesis, metabolism, and RNA processing in the nasal mucosa. In lung tissue, pathways overlapped with the nasal mucosa but also included lung-specific enrichment in extracellular matrix organisation, and more apoptosis pathways were enriched in the lung tissue. Overall, the lung displayed a higher number of regulated substrates (47 vs 42), with several of them shared. This suggests that the infection induces more pronounced cell death and tissue repair/fibrotic responses in the lower respiratory tract.

Given the predominance of arginine (R) at the P1 position across both nasal mucosa and lung tissues, we next focused on R-specific cleavage events to delineate their protease contributors and functional implications in tissue-specific proteolytic regulation. In the infected nasal mucosa and lung tissue most of these cleavages were presumably made by mainly KLKB1, CTSB. FURIN and ST14.

We observed prominent cleavage following N in infected lung tissue, which was also among the top cleavages in nasal mucosa, likely mediated by LGMN. LGMN is primarily a lysosomal protease involved in protein degradation and antigen processing but can also be secreted. It is known to activate pro-cathepsins that regulate ECM remodelling, many of which are strongly modulated in both infected tissues. Across the infected nasal mucosa tissue, we identified 15 putative LGMN substrates that were regulated in the global proteome: 9 increased, associated with metabolism, protein translation, and immune signalling, and 6 decreased, linked to immunity, metabolism, and apoptosis. In the lung, 10 substrates were observed in the global proteome, of which 2 increased (also increased in nasal mucosa) and were involved in immune processes, while the remaining 8 decreased, affecting immunity as well as metabolism and apoptotic pathways. These substrate-specific changes reflect a dual response: promotion of tissue remodelling and immune signalling, alongside suppression of translation, protein turnover, and cellular homeostasis. Together with broader proteomic analyses showing coordinated cleavage of upregulated proteins to modulate signalling and ECM, and of downregulated proteins to restrain innate immunity, antigen presentation, and intracellular defences, these data highlight LGMN as a central regulator orchestrating tissue-specific remodelling, host defence, and stress responses during infection.

In the nasal mucosa, cleavage sites showed a preference for G, a pattern consistent with activity from the broad-specificity proteases CAPN1/2 as well as ELANE. In the global proteome, HBB and FN1 were increased, while ACTN4 was decreased, and all three are presumed to be cleaved by either the calpains or ELANE.

Overall, the analysis of infected nasal mucosa and lung tissues revealed a coordinated, protease-driven remodelling program affecting multiple pathways, including core immune and cellular processes and apoptosis. Although both tissues share some of the same substrates, cleavage patterns differ, the nasal mucosa exhibits broader but primarily Arg-, Gly-, and Asn-directed cleavage, affecting multiple pathways, whereas the lung showed selective Asn- and Arg-directed proteolysis, focusing on ECM remodelling and targeted intracellular inactivation. Furthermore, the lung exhibiting more regulated substrates and stronger apoptotic and ECM remodelling signatures. LGMN emerged as a key modulator of many of the processes, linking proteolytic regulation to antigen processing, ECM remodelling, and coordinated control of immune and metabolic pathways. In contrast, CAPN1/2 and ELANE appear to shape nasal responses through cleavage of cytoskeletal and structural proteins. Together, these findings highlight site-specific protease activities driving differential tissue remodelling, host defence, and homeostatic regulation across the upper and lower respiratory tract during infection.

Global proteome and N-terminomics analyses identified 112 proteases, however, these methods do not discriminate between inactive and active forms. We therefore applied targeted proteomics to the nasal mucosa and lung, where most proteases were quantified, to confirm their presence and assess their active abundance.

### Differential protease cleavage programs in nasal and lung tissues during influenza A infection

To confirm the protease activity observed and measure the active protease fraction directly, we performed targeted proteomics analysis (PRM) on the nasal mucosa and lung tissue. Based on the cleavage pattern observed by the N-terminomics analysis, we chose 30 of the 112 proteases for further validation. In addition, four proteases involved in HA activation (Furin, PCSK6, TMPRSS2, and TMPRSS4) were also included. All selected targets were incorporated into the spectral library and are listed in Supplementary Table S1. For this validation, we created a targeted proteomics PRM assay to monitor protease identification. We designed an assay with 121 peptides, spanning quantification and activation peptides. We created a spectral library with the heavy peptides to match the elution and fragmentation profile of the endogenous peptides. We were able to quantify 28 of the proteases by using 47 the heavy peptides as a guide for finding the correct peak for the light/endogenous peptides (Fig. 3A). The protein expression profile of all peptides can be seen in Supplementary Table S11 and Supplementary Figures S3).

**Figure 3.**
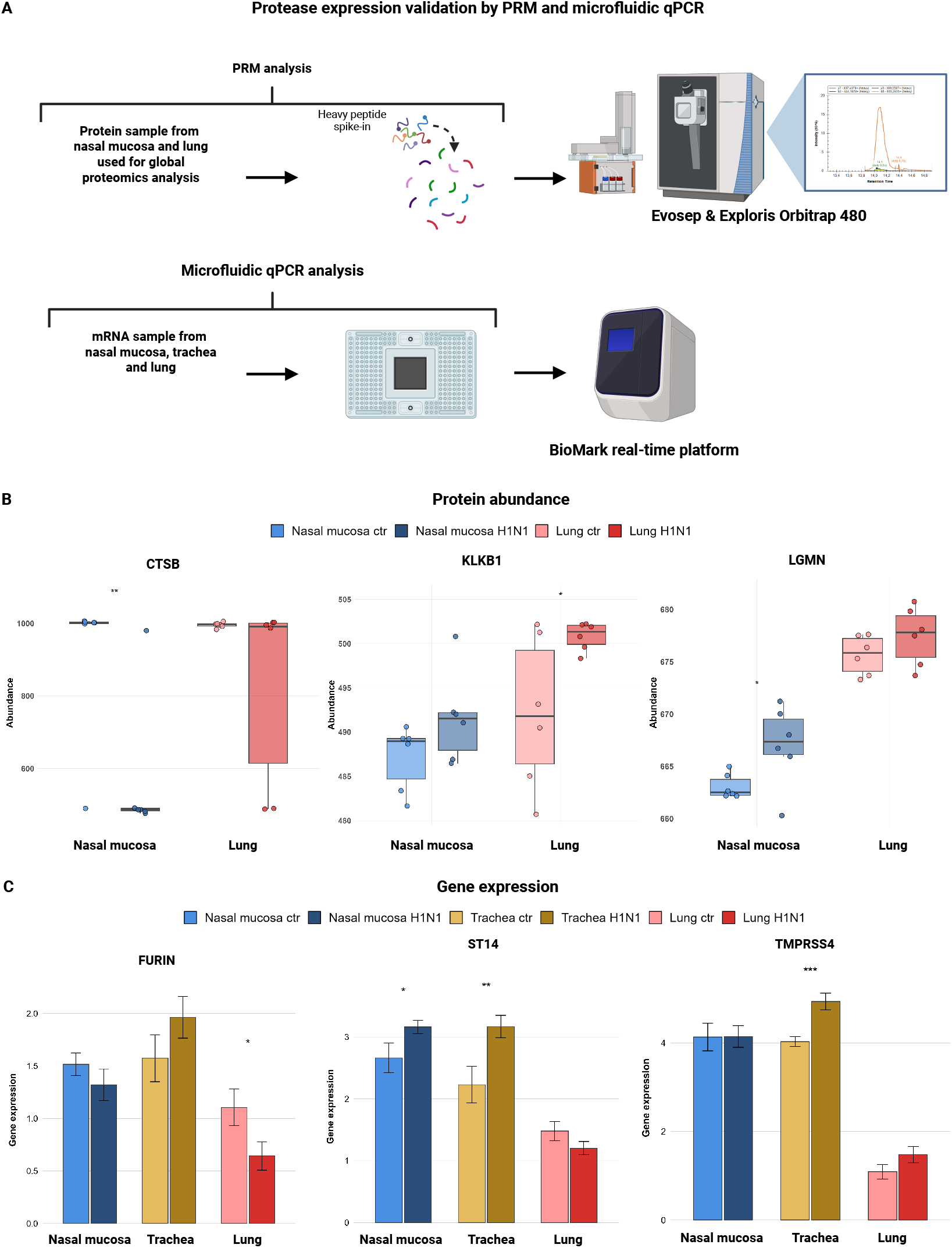
Validation of protease expression across respiratory tissues. A. Experimental design of two validation approaches, one by targeted proteomics (PRM) using the nasal mucosa tissue and the lung. Another by microfluidic qPCR using nasal mucosa, trachea and lung tissue, created in Biorender. B: Protein expression by PRM, blue colour is the nasal mucosa, and red colours are the lung, for CTSB, KLKB1 and LGMN. C: Gene expression by microfluidic qPCR, blue colours are the nasal mucosa, yellow colours are the trachea, and red colours are the lung for Furin, ST14 and TMPRSS4. Statistical significance is indicated as follows: p < 0.1 (*), p < 0.05 (**) and p < 0.01 (***).

We additionally profiled transcripts for fifteen proteases by microfluidic qPCR (Supplementary Table S13) in upper and lower respiratory tissues to provide complementary information to the proteomics data. Several proteases were found significantly regulated in all tissues upon infection, including TMPRSS4, Furin and ST14, which are all known to facilitate HA cleavage (Fig. 3C) (Supplementary Table S14 and Supplementary Figures S4).

Confirming our prior observations, ten of the proteases were seen significantly abundant in either the nasal mucosa tissue or the lung tissue between the groups in the PRM analysis (Supplementary Table S11 and Supplementary Figures S3). Some proteins showed significant changes in PRM analysis that were not detected in global proteomics, reflecting PRM’s higher sensitivity and ability to quantify low-abundance or isoform-specific peptides.

In the nasal mucosa, ACE, CTSB, and F2 proteins decreased after H1N1 infection, whereas METAP2, PSMB9, and LGMN increased (Fig. 3B and Supplementary Figures S3), reflecting enhanced antigen processing and lysosomal proteolysis. In the lung, KLKB1 and ST14 increased while PLG and PRSS8 decreased, highlighting tissue-specific proteolytic programs that balance inflammation, ECM remodelling, and tissue integrity during infection (Fig. 3B and Supplementary Figures S3).

Using the established library, we confirmed the active fraction of five proteases using 18 activation peptides (Fig. 4A). Based on the data on the active and inactive (spanning) peptide, we assessed and validated the activity of CTSD and LGMN proteases in the nasal mucosa and lung tissue, observing differences between infected and healthy samples. However, it is difficult to make a clear conclusion on CTSC, PLG and PRSS8 as we only observed the active part of the protease, not the inactive (spanning peptide) (Supplementary S16).

**Figure 4.**
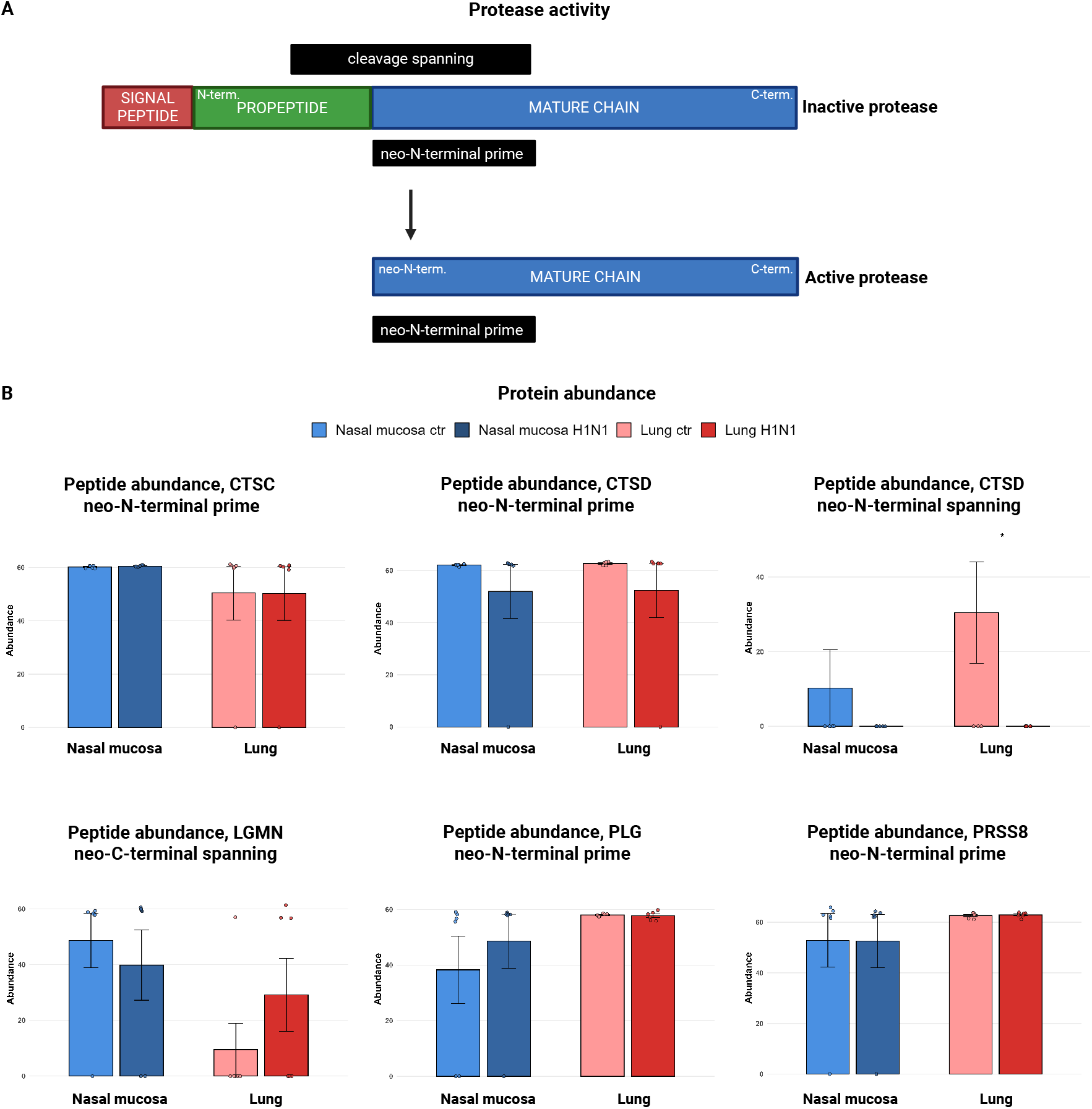
Activity of proteases. A. Schematic of protease activity once propeptide is cleaved. B. Peptide expression by PRM, blue colours are the nasal mucosa, and red colours are the lung. CTSC neo-N-terminal prime. CTSD neo-N-terminal prime. CTSD neo-N-terminal spanning. LGMN neo-C-terminal spanning. PLG neo-N-terminal prime. PRSS8 neo-N-terminal prime. Statistical significance is indicated as follows: p < 0.1 (*), p < 0.05 (**) and p < 0.01 (***).

In the nasal mucosa, no significant difference was observed between the ctr and the H1N1 infected tissue between any of the peptides. However, there was a decrease in abundance in the CTSD neo-N-terminal spanning peptide between the H1N1 infected and the control samples (Fig. 4B). The same was observed in the lung tissue, where the CTSD neo-N-terminal spanning was significantly decreased in the H1N1 infected lung tissue compared to the control (Fig. 4B), indicating more active CTSD in the infected nasal mucosa and lung tissue compared to their respective controls. An increase of the LGMN neo-C-terminal spanning peptide in the H1N1 infected lung tissue indicates inactive LGMN in the H1N1 infected lung tissue, which also confirms the results from the enriched samples (Fig. 4B).

Although we were unable to monitor a spanning peptide for CTSC, PLG or PRSS8, the neo-N-terminal peptide of CTSC confirmed that the protease is active in both the nasal mucosa and lung, consistent with observations from the enriched samples. For PLG and PRSS8, cleavage at the propeptide in the enriched samples indicated that it is active. However, no significant differences were observed between the two conditions in either tissue. PRM analysis corroborated these findings, confirming that PRSS8 is active in both viral infected and control tissues.

In summary, targeted PRM analysis validated key findings from global proteomics and revealed additional changes not captured in untargeted data. Microfluidic qPCR further confirmed several of these alterations at the transcriptional level, highlighting the sensitivity and specificity of the targeted approach for detecting tissue- and infection-associated protease regulation. Assessment of select proteases confirmed their activity in both nasal mucosa and lung tissue, consistent with observations from the enriched samples.

## Discussion

In this study, we profiled the proteomic response along with the protease network in the nasal mucosa, trachea, and lung of pigs three days post-H1N1 IAV infection. We identified 112 proteases in the three respiratory tissues, corresponding to nearly 20% of the more than 600 protease-encoding genes reported in in humans^19^, highlighting that the respiratory tract expresses a substantial fraction of the human protease repertoire. Of these, 28% were serine proteases, 29% cysteine proteases, 21% metalloproteases, and 22% belonged to other classes. We confirmed protease changes using microfluidic qPCR and targeted proteomics (PRM).

None of the 112 proteases identified showed significant abundance changes in the trachea in either the global proteome or the enriched samples, whereas several were differentially abundant in the nasal mucosa and lower lung tissues upon viral infection. Some of the identified proteases have previously been linked to IAV infection. However, most existing studies were conducted in human cell lines, patient samples, or in other animal models such as mice and ferrets^6–9,11,20–22^. Several studies in pigs have focused on a few specific proteases, including TMPRSS11D, TMPRSS2, and FURIN^23–25^. Here, we extend these efforts by employing an N-terminomics approach to investigate host proteases in respiratory tissues following IAV infection in a porcine model.

H1N1 infection in pigs elicited a classical antiviral innate immune response across the respiratory tract. Antiviral proteins were increased, particularly in the nasal mucosa and trachea, while influenza A and NOD-like receptor signalling pathways were enriched. The nasal mucosa showed the highest number of differentially expressed proteins, indicating it as the primary site of viral replication and immune activation. In contrast, the lung exhibited, lower viral load, but severe tissue damage, suggesting immune-mediated pathology rather than active viral replication, which correlates with previous findings^26^. Notably, no viral proteins were detected, despite searching against reference proteomes containing H1N1 sequences, which may reflect the complexity of tissue proteomes and limited viral protein abundance. Overall, our results suggest a classical antiviral response in the upper airways, consistent with observations by Laybourn et al., who reported that the mxH1N1 strain elicited the strongest and most prolonged antiviral response at 4 days post-infection compared to human- and swine-adapted strains^27^. Our results reflect a similarly robust innate immune activation in the upper respiratory tract.

Several proteases previously implicated in influenza virus activation, including TMPRSS11D, TMPRSS4, FURIN, and ST14, showed tissue-dependent differences in abundance, suggesting that the potential for HA cleavage and viral spread varies along the respiratory tract. In contrast, TMPRSS2, a well-characterized HA-activating protease in humans^6^, remained unchanged after infection, indicating that other proteases may play more prominent roles in the porcine airway tissue specific response. These observations imply that host proteolytic capacity, and consequently viral replication dynamics, are modulated in a tissue-specific manner, providing insight into how protease distribution may influence infection outcomes and identify potential targets for antiviral intervention.

In the uninfected pigs, the lung exhibited a more extensive and active proteolytic environment than the upper airways, consistent with its complex immune and metabolic demands. Although individual proteases and inhibitors showed tissue-specific enrichment, the overall cleavage motif patterns in healthy lung and nasal mucosa closely resembled those observed during infection. This similarity suggests that baseline proteolytic activity is already configured to support key physiological processes and may prime tissues for rapid remodelling and immune activation upon viral challenge.

Following infection, the nasal mucosa exhibited pronounced activation of the proteolytic network, reflecting its role as the primary interface with inhaled pathogens. Proteases with both broad and narrow substrate specificity were increased, supporting both direct antiviral activity and the processing of signalling molecules involved in immune responses. This expanded protease activity occurs alongside a relatively high viral load and elevated levels of antiviral proteins in the nasal mucosa. These observations suggest a dynamic tissue environment in which broad protease activity could influence both pathogen processing and local host responses. Whereas the infected lung tissue exhibited a more distinct cleavage pattern related to inflammation but also ECM remodelling, linking the protease abundance to the lung pathology observed in a previous study, where the mxH1N1 infection resulted in severe lesions in the lower lung tissue^26^.

A common observation across both nasal mucosa and the lung was the preference for arginine (R) at the P1 position. In the nasal mucosa, we observed increased levels of several proteases with a strong specificity for arginine at P1, including DPP3, FURIN, KLKB1, and PCSK6. Notably, substrates of CTSB were differentially regulated despite no significant change in overall CTSB abundance between conditions. A similar pattern was observed in lung tissue: substrates of CTSB showed clear regulation, yet the proteases most strongly regulated with a preference for arginine at P1 in the lung were F2, KLKB1, PCSK6, and ST14. These findings suggest that tissue-specific regulation of protease activity, rather than protease abundance alone, shapes substrate processing preferences.

The broad cleavage pattern observed in the nasal mucosa, which includes amino acids such as glycine (G) and alanine (A), may be driven by the increased abundance of proteases, including DPP7, CTSD, and ELANE^16^. Notably, CTSD activity could be directly assessed, revealing higher levels of active enzyme within infected tissues. Given its involvement in pathways such as ECM remodelling, antigen processing, and apoptosis, the upregulation of CTSD may reflect a dual role in host defence and tissue pathology. Specifically, intracellular CTSD could facilitate antigen processing and immune signalling, while excessive activity may promote apoptosis and ECM degradation^28,29^, potentially contributing to local inflammatory responses during viral infection.

Another common cleavage preference shared between the two infected tissues was asparagine (N), which was particularly predominant in the lung tissue. This preference may reflect increased activity of LGMN, which was elevated in both the infected lung and nasal mucosa. LGMN participates in multiple pathways, including protein degradation, antigen processing, inflammatory signalling, and ECM remodelling^30^. By targeted proteomics, we monitored the neo-N-terminal spanning peptide of LGMN but did not observe a difference between conditions despite the overall increase in total protein abundance in the infected tissues. This could indicate that a fraction of active LGMN is secreted or redistributed, potentially contributing to ECM remodelling or influencing viral entry, although the precise extracellular role during infection remains to be clarified. While the role of LGMN in IAV has not been described previously, a recent study by Dinesh et al. identified LGMN as a host protease that contributes to SARS-CoV-2 entry in an ACE2-dependent manner^31^. Together, these observations suggest that it would be valuable to investigate further the mechanism of LGMN in relation to IAV infection.

In conclusion, our study reveals distinct, tissue-specific proteolytic responses in the porcine respiratory tract during H1N1 infection. The nasal mucosa exhibits robust protease activation and antiviral responses, whereas the lung displays a unique cleavage pattern linked to inflammation and extracellular matrix remodelling, consistent with tissue pathology.

## Methods

### Ethical approval declarations

The animal experiment was performed under biosafety level 2 conditions and an animal study protocol approved by the Danish Animal Experimentation Council (protocol no. 2020-15-0201-00502).

### Preparation of virus inoculum

The H1N1 strain, A/Swine/Mexico/AVX-39/2012, kindly provided by Nacho Mena and Adolfo Garcia-Sastre of Mount Sinai School of Medicine was propagated and passaged three times in Madin–Darby canine kidney (MDCK) cells^26^. The viruses were stored at -80°C before inoculation. The titers were determined by tissue culture infectious dose 50% (TCID50) assay in MDCK cells and diluted in Eagle’s Minimum Essential Medium (Gibco) to a TCID_50_/mL of 10^7^. The IAV isolate was selected to resemble a pre-pandemic variant of the 2009 pandemic influenza A virus subtype H1N1^32^.

### Experimental design

The experimental setup has previously been described in detail, however, only a subset of animals described in that paper were included in the present study^26^. The experiment included 12 confirmed IAV-negative, seven-week-old Danish Landrace Crossbred pigs. The pigs were allocated into two groups and housed in separate isolation units. They were acclimatised for one week, fed non-pelleted feed, and had ad libitum access to water. Group 1 (control) and Group 2 (infected) included six pigs each. The pigs were sedated before inoculation. Group 1 was mock inoculated intranasally in one nostril by a MAD nasal device (Teleflex) containing 3 mL culture medium only, while group 2 was inoculated with 3 mL 10^7^ TCID_50_/mL of a H1N1 strain, A/Swine/Mexico/AVX-39/2012. The pigs were euthanised 3 days post inoculation, and tracheal, nasal mucosa, and lower lung tissue (*lobus dexter caudalis*) were collected and washed in PBS, snap frozen in liquid nitrogen, and stored at -80°C for proteomics analysis. Nasal mucosa, trachea, and lower lung (*lobus dexter caudalis*) tissues were also collected for mRNA extraction, these tissues were stored in RNAlater at -20°C.

### Protein extraction

All reagents are from Sigma Aldrich unless otherwise specified. The mucosal membranes of trachea and nasal tissue were stripped from the cartilage, and all three tissues were homogenized in lysis buffer (6 M guanidine hydrochloride (GuHCl), 10 mM tris(2-carboxyethyl)phosphine (TCEP), 40 mM chloroacetamide (CAA), 50 mM 4-(2-hydroxyethyl)-1-piperazineethanesulfonic acid (HEPES), pH 8.5, with cOmplete^™^ Mini EDTA-free protease inhibitor cocktail) using a TissueLyser II (Agilent) and one steel bead twice for 1 minute at 1–30 Hz. The samples were incubated for 5 minutes at 95°C for reduction and alkylation of cysteines. To remove contaminants, lysates were precipitated in acetone overnight at -20°C, centrifuged for 20 minutes at 4000x g at 4°C, and resuspended in buffer solution (2.5 M GuHCl, 520 mM HEPES, pH 8.5). Protein concentration was determined by NanoDrop One (Thermo Fischer Scientific).

### HUNTER DIA

All reagents are from Sigma Aldrich unless otherwise specified. Protein extracts from nasal mucosa, trachea and lung from six control animals and six H1N1 infected animals were used for HUNTER^33^. 50 µg of extracted protein from each sample was diluted in TAILS buffer (2.5 M GuHCl, 520 mM HEPES, pH 8.5) to 100 µL total volume. 60 mM of formaldehyde and 50 mM of sodium cyanoborohydride were added to each sample. The pH was adjusted to 6-7 with 2 M HCl and incubated at 50°C for 1.5 hours, 450 rpm. After 1.5 hours, the samples were vortexed and spun down, and 100% ethanol was added to each sample to achieve a final ethanol concentration of 80%.

Sera-Mag SpeedBead Carboxylate-Modified Magnetic Particles (SP3 beads) were prepared according to the manufacturer’s instructions. 1:10 ratio protein:beads w/w were prepared by pipetting gently up and down, the beads were washed three times with MiliQ water, the beads were added to each sample and incubated at room temperature for 15 minutes at 800 rpm for binding. The tubes were then placed on a magnetic rack (Thermo Scientific), and the supernatant was removed. The beads were gently resuspended in 90% ethanol for 0.5 µg/µL protein concentration in each sample, assuming no loss. The tubes were placed back on the magnetic rack, and the supernatant was removed. The washing steps were repeated a total of three times. The beads were resuspended in 100 mM HEPES for a final concentration of 1 µg/µL. The proteins were digested overnight with trypsin (Promega) at a ratio of 1:50 trypsin to protein (w/w) at 37°C at 350 rpm. The samples were vortexed and spun down before adding 1:1 volume of 100mM HEPES to each sample. Each sample was sonicated for 1 minute and placed back on the magnetic rack for 2 minutes. The supernatant was transferred to new tubes after binding, and the beads were discarded. 10% of the digestion solution was set aside as a non-pull out (NPO) sample from each of the samples and stored at -20°C. 100% ethanol was added to the remaining solution for a final concentration of 40% ethanol and mixed. Undecanal was added to each sample 50:1 undecanal to peptide (w/w) ratio, along with 5 M sodium cyanoborohydride (NaBH_3_CN) in 1 M NaOH for a final concentration of 50 mM NaBH_3_CN, and the pH was adjusted to be around 6-7 with 2 M HCL before the samples were incubated at 50°C for 1.5 hours. The samples were vortexed and were acidified to pH 2 using buffer H’ (40% ethanol, 5% TFA). The volume was normalised to 200 µL using buffer H (40% ethanol, 1% TFA). Each pull-out (PO) sample was desalted using a Solaµ plate. The filters were conditioned using 200 µL 100% methanol and spun down for 1 minute at 1500 rpm. 200 µl of buffer H was added to each column and then spun down for 1 minute at 1500 rpm. This step was repeated before loading each sample to the columns. The flow-through contained the PO sample, each sample was dried in a SpeedVac for 30 minutes at 45°C and stored at -20°C. 500 ng of the PO and NPO samples were loaded on Evotips according to the manufacturer’s loading protocol.

For liquid chromatography/mass spectrometry (LC-MS) analysis, an Evosep One HPLC system was used in-line with an Orbitrap Exploris 480 mass spectrometer (Thermo Fischer Scientific). The instrument was operated in positive polarity in data-independent acquisition (DIA) mode, with a gradient of 58 minutes using an Aurora TS column (15 cm length, 75 µm diameter, and bead size 1.7 µm) running at 100 nl/min, with column heating at 50°C, and a transfer tube temperature of 240°C.

Global parameters were set to nano-spray ionisation, with a static positive ion voltage of 2000V and 600V for the negative ion. The resolution of Orbitrap for MS/MS scan acquisition was set to 120,000 with FAIMS ON with –45 CV at standard resolution mode. The RF lens for both MS and MS/MS was 40%.

The normalised AGC target was set to 300%. A scan range of 400-1350 m/z for the MS scan, and the maximum injection time was automatic. The Orbitrap resolution for the DIA scans was 150,000. The normalised AGC Target was 1,000% for the MS/MS DIA scans, and the maximum injection time was automatic. The scan range for the MS scans was 400–1350 m/z. The scan range for the MS/MS DIA scans was 400-1000 m/z. The HCD collision energy was 33%. The window size of the DIA scan was 20 m/z, with an overlap of 1 m/z, and 29 events per scan.

### Targeted proteomics

After screening by HUNTER, 34 proteases were chosen for further validation, based on the results obtained from the screening (Supplementary Table S1). Quantification peptides were designed based on a list of criteria: First, the peptides must be unique for the protease in question, and the peptides must be located in the activated form of the enzyme. Their length should ideally be between eight and twenty amino acid residues, and peptides containing methionine residues and/or missed cleavages were avoided. Using these criteria, we selected the best three peptides per protease to establish our targeted proteomics assay. However, a lot of the selected porcine proteases were not well annotated, for those where the sequence from sus scrofa was too diverse from the human sequence, PeptideCutter was used to locate sequences for quantification.

When it comes to activation peptides used for the targeted assay, they were selected from the region spanning the cleavage site or from the right flank at either the N- or C-terminal cleavage site of the propeptide, according to the specific activation mechanism of each target protein. They were between six and thirty amino acids in length, and were cleavable by either trypsin or LysC. Based on these criteria, we obtained 121 heavy peptides (Heavy arginine (Arg U13C6;U-15N4), heavy lysine (Lys U-13C6;U-15N2), carbamidomethylation of cysteines) to be used for quantification and monitoring activation of proteases, synthesised (JPT Peptide Technologies, Berlin, Germany) (Supplementary Table S2). Using these peptide sequences, we created a spectral library of the proteases. We injected 7 fmol per peptide by 10^5^ dilution of stock.

For liquid chromatography/mass spectrometry (LC-MS) analysis, an Evosep One HPLC system was used in-line with an Orbitrap Exploris 480 mass spectrometer (Thermo Fischer Scientific). The instrument was operated in positive polarity in data dependent acquisition (DDA) mode, with a gradient of 31 minutes using an Aurora TS column (15 cm length, 75 µm diameter, and bead size 1.7 µm), running at 100 nl/min, with column heating at 50°C, and a transfer tube temperature of 240°C. Full MS scans (MS1) were acquired at a resolution of 60,000 over a mass range of 375–1500 m/z, with an AGC target of 300, and a maximum injection time was set to Auto. Data-dependent MS/MS scans (MS2) were performed using HCD fragmentation at 30% normalised collision energy, with a resolution of 30,000, an AGC target of 100, and a maximum injection time set to Auto.

For the PRM assay, a Skyline document was created with the peptide sequences. These peptides were used for the inclusion list that was exported and used for the targeted method settings.

The protein extracts from nasal mucosa and lung from six control animal and six H1N1 infected animals was used for PRM analysis. A total peptide amount of 500 ng for each sample was loaded on EvoTips according to manufacturer instructions. Briefly, the tips were immersed in isopropanol for 30 sec, buffer B 20 µl centrifugation for 1 min at 700 g, buffer A conditioning, load sample, buffer A washing, and store in buffer A 120 µl. Heavy isotope–labeled synthetic peptides (JPT; 0.7 nmol per peptide aliquot) were diluted 1:100,000 and spiked into each sample as internal standards prior to analysis. The diluted heavy peptide mixture was added at a volume of 3.7 µL per sample. The PRM data were acquired on an Orbitrap Exploris 480 mass spectrometer with an Evosep One HPLC system, with the same setup as the previous run. Targeted precursor ions were isolated with an isolation window based on the transition list, which was specific for each peptide. Full MS scans were acquired at 60,000 resolution, with an AGC target of 300 and a maximum injection time set to Auto. Scan range was set to 375-1500. RF lens was 40% and the mass tolerance was 10 for high and low. PRM scans of the selected precursors were acquired at 15,000 resolution.

### Quantification of IAV and viral titration

RNA extraction for quantification of IAV was performed as described previously^26^.

### Microfluidic qPCR

Sixteen proteases were chosen for validation by microfluidic qPCR, along with six potential reference genes. Primers for the selected proteases are shown in the Supplementary Table S3. Primer design, cDNA synthesis, and cDNA preamplification were performed as described before^27^. All twelve of each tissue including: nasal mucosa, trachea, and lower lung samples were used for the microfluidic qPCR (6x control and 6x H1N1 infected). Microfluidic qPCR was carried out using 96.96 Dynamic Array IFC chips (Standard BioTools Inc.) on the BioMark real-time platform using a previously described PCR protocol^27^.

### Data and Statistical Analysis of HUNTER DIA

The raw files generated were used to screen and select candidates for targeted proteomics experiments. The data was analysed using Spectronaut (19.1.240806, Biognosys AG). The acquired spectra were searched using the directDIA option in Spectronaut. Searches were run using Precursor PEP cutoff of 0.1, Protein Qvalue cutoff (run) of 0.05, and Protein PEP cutoff of 0.5. The minimum length of the peptide was set to 6, and the maximum length was set to 46. Carbamidomethyl (C), DimethLys0 were set as fixed modifications, and Acetyl (protein N-term), Deamidation (N), DimethNter0 and Oxidation (M) were set as variable modifications. Default settings were used for tolerances, and 2 missed cleavages were allowed. Quantification was performed using Spectronaut’s version of MaxLFQ algorithm with precursor quantification on MS1 level. No cross-run normalisation and no imputation method were used. We used the Sus scrofa reference proteome database from UniProt^18^ (Downloaded October 2024).

The statistical analysis was conducted on the protein and peptide level, using R Studio (R version 4.5.0), using the following packages: tidyverse (2.0.0), readxl (1.4.5), ggplot2 (4.0.0), dplyr (1.1.4), purr (1.1.0), readr (2.1.5), tidyr (1.3.1), broom (1.0.10), stringr (1.5.2), ggVennDiagram (1.5.4), clusterProfiler (4.16.0), AnnotationDbi (1.70.0) and org.Ss.eg.db (3.21.0).

The protein data was imputed, normalised based on the median for each channel, and log2 transformed. The NPO samples were extracted from the protein data, and a two-way ANOVA test, followed by a Tukey post hoc analysis to correct for multiple testing, was done on the protein level. Differentially expressed proteins (DEPs) were defined with a p-adjusted value (padj) below 0.05 and with a log2 fold change (lfc) of ±1. Exploratory functional enrichment analysis of detected protein groups was performed using Gene Set Enrichment Analysis (GSEA) against the Kyoto Encyclopedia of Genes and Genomes (KEGG) and Gene Ontology (GO) databases. Ranked gene lists, based on log2 fold-change values, were analysed using the gseKEGG and gseGO functions from the clusterProfiler package. All genes were included in the analysis (pvalueCutoff = 1) to allow comprehensive exploration of potentially enriched pathways and GO terms, with pathway sizes restricted to a minimum of 5 and a maximum of 500 genes.

The PO samples were extracted from the peptide data, and the data were normalised. First, the median from each channel of the NPO samples was used to normalise the PO samples. The data was log2 transformed, and this data was used for absolute peptide level data analysis. Secondly, the peptide data was normalised based on the abundance of the parent protein to better capture cleavage dynamics resulting from altered substrate concentration versus protease activity.

The dataset was annotated in Python (version 3.9) using CLIPPER 2.0^34^ which was used to predict proteases and protease inhibitors in each sample. Following the annotation, statistical analyses were carried out in RStudio, where a two-way ANOVA was performed, followed by a Tukey post hoc test to assess differences in the abundance of predicted proteases and inhibitors across tissue types. The MEROPS database^16^ used to investigate each protease’s known substrates and their cleavage patterns. The list of substrates for each protease was used to determine whether these substrates were differentially abundant and cleaved in the different tissues. The substrate cleavage environments (residues P1-P1’) were used to create ice logos using Seq2Logo 2.0^35^, using default settings. Significantly abundant substrates (padj > 0.1) were annotated using Reactome (ReactomePA version 1.52.0) to determine their associated pathways in R studio, the human and pig proteomes were both used for this analysis.

### Data and statistical analysis of targeted proteomics

The heavy peptide injections were used to generate a library, the samples were analysed using Proteome Discoverer (version 2.4.0.305, Thermo Fisher Scientific) and the library was generated using Skyline21 (version 25.1.0.142). Out of the 121 peptides spiked, 93 peptides were identified and used to generate a transition list for the targeted analysis. Each sample analysis by the Exploris Orbitrap 480 was analysed in Skyline as well, using the library generated previously to select the appropriate ions for quantification. The analysis in Skyline was curated manually by filtration according to the heavy peptide elution and fragmentation profile. A range of 3 to 6 ions has been used for the quantification. Two peptides were used to assess protease activity: A spanning peptide, and depending on the position of the propeptide, either a C-terminal–right or N-terminal–right peptide was employed to determine whether the protease was active. No normalisation was done on the quantification dataset, whereas the activation dataset was normalised based on the total protein abundance for each protease from the quantification dataset. The transition result report was analysed using R Studio, and a Welch two-sided t-test was conducted between the control and H1N1 infected group. This was done to determine whether the specific proteases were significantly abundant in their respective tissue, the peptide and protein expressions were considered significant with a p-value below 0.1 and lfc of ±1.

### Data and statistical analysis of microfluidic qPCR

Data was processed as described previously^27^. The statistical analysis was conducted using R Studio (R version 4.5.0). A biologically relevant cut-off value of lfc of ±1 compared to the control was used. The change in gene expression level was considered statistically significant with a p-value below 0.1 (Student’s t-test) and lfc value of ±1. A Pearson correlation was conducted to determine the correlation between protein and mRNA using R Studio.

## Supplementary information

Supplementary data supporting the publication can be found on Zenodo with doi: doi.org/10.5281/zenodo.18591198. Supplementary Figures S1, S2, S3 and S4 are included at the end of this document.

## Data availability

The proteomics and degradomics dataset have been deposited to the ProteomeXchange Consortium via the PRIDE^36^ partner repository with the dataset identifier PXD074287. Reviewers can access the dataset with username and password s2C70klUCspp. The targeted proteomics experiment files have been deposited to PanoramaWeb and can be accessed with the link https://panoramaweb.org/4W3hr8.url with reviewer and password +GCS$qnW!1y2Pr.

## Acknowledgements

This work was supported by the Novo Nordisk Fonden (NNF19OC0056326 to L.E.L). K.K. is supported by a Novo Nordisk Foundation Young Investigator Award (grant no. NNF16OC0020670) and a postdoctoral fellowship grant from the Independent Research Fund Denmark (grant no. 4257-00010B). The funders had no role in study design, data collection and analysis, decision to publish, or preparation of the manuscript. We are grateful to the DTU Proteomics Core Facility for maintenance and running of instruments. We especially thank M. Wennekers Nielsen and M. Vestegaard Lukassen for their help and advice during troubleshooting, sample preparation and data acquisition. We thank Karin Tarp for her technical assistance.

## Author contributions statement

K.S., U.a.d.K., K.K., and C.H.P. conceived and designed the research. C.H.P. and A.M.H. performed sample preparation and mass spectrometry analysis. C.H.P. prepared and analysed the datasets with feedback from K.S. and K.K. K.S. analysed the micro qPCR data, and C.H.P. performed additional data analysis. C.H.P. wrote the manuscript with input from all authors. All authors read and approved the final version of the manuscript.

## Supplementary figures

**Supplementary Figure S1.**
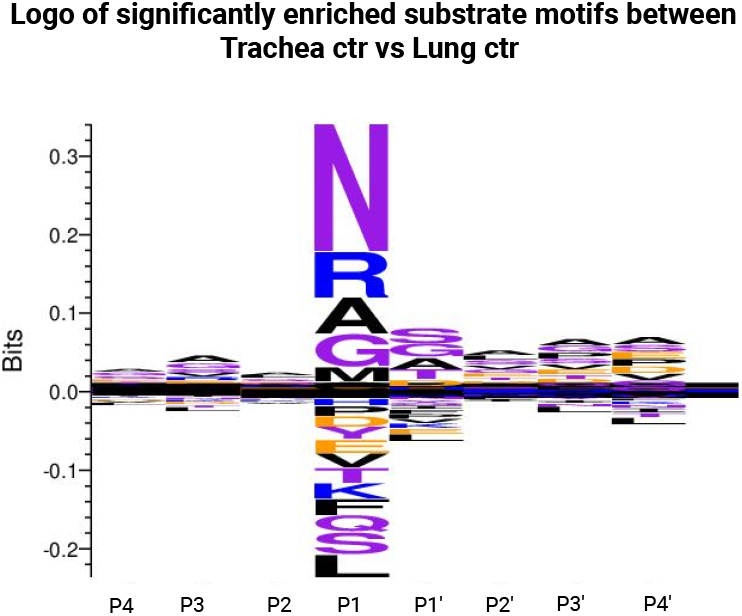
Sequence logos

**Supplementary Figure S2.**
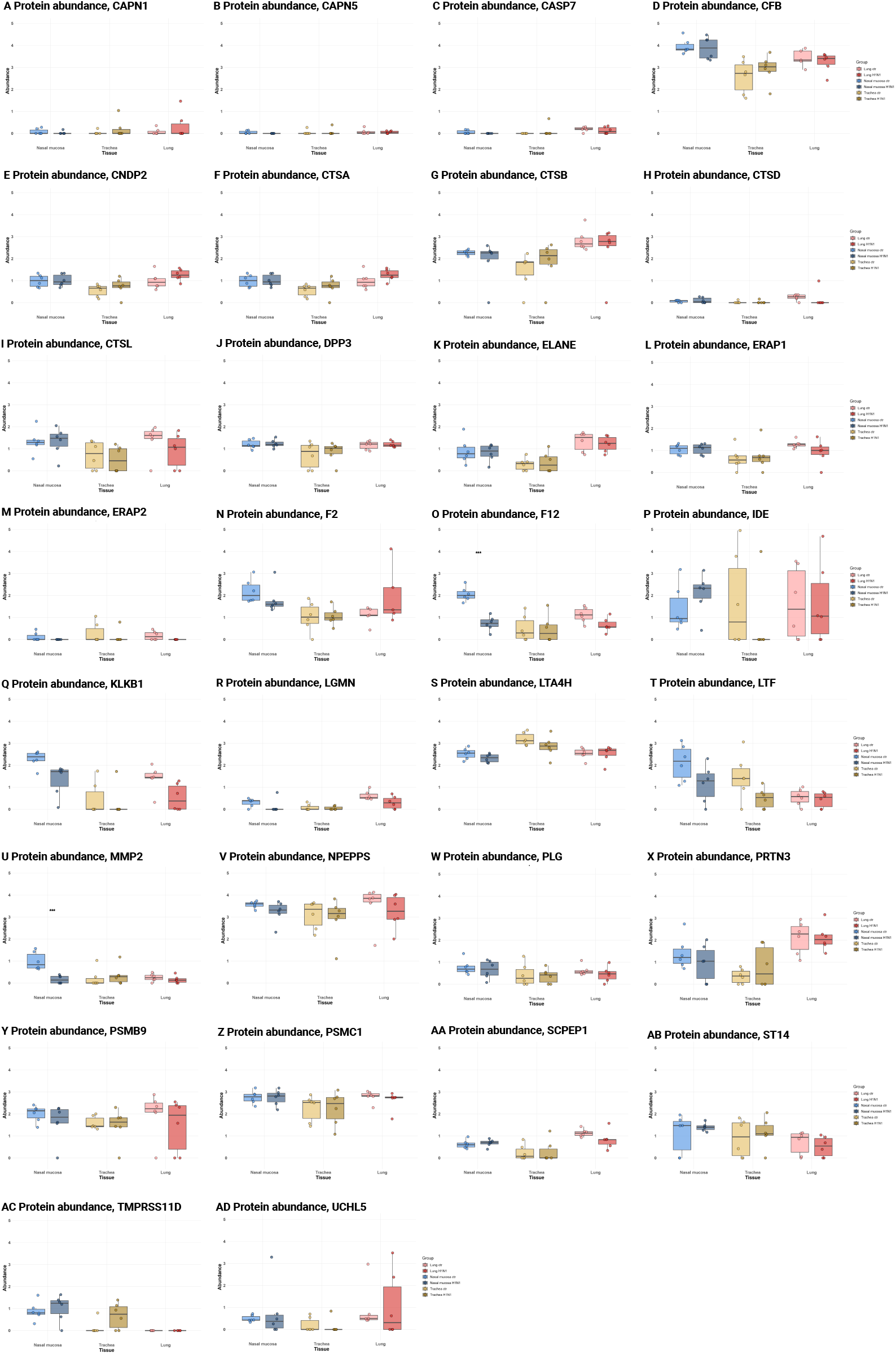
Protease abundance, global proteomics

**Supplementary Figure S3.**
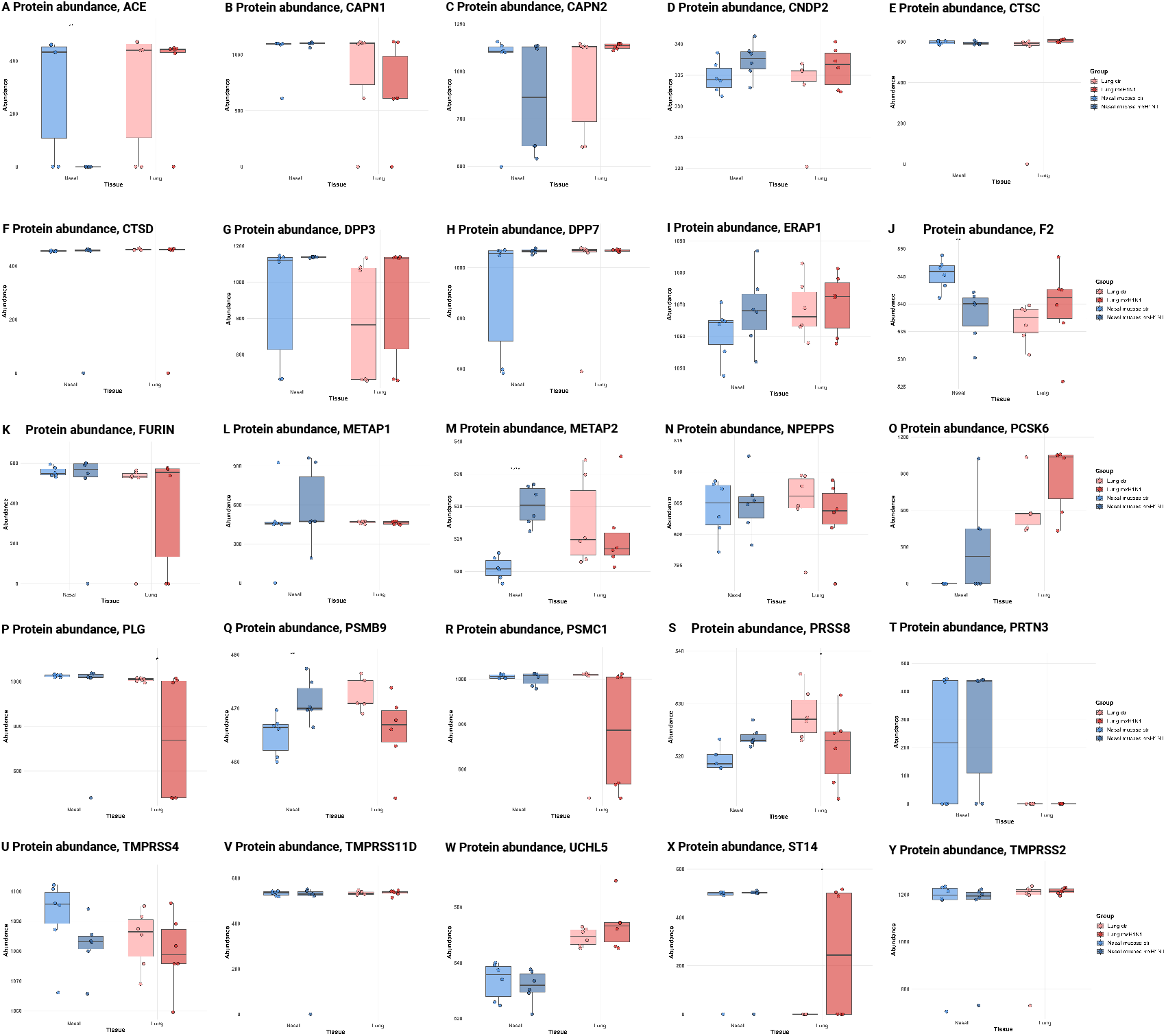
Protease abundance, targeted proteomics

**Supplementary Figure S4.**
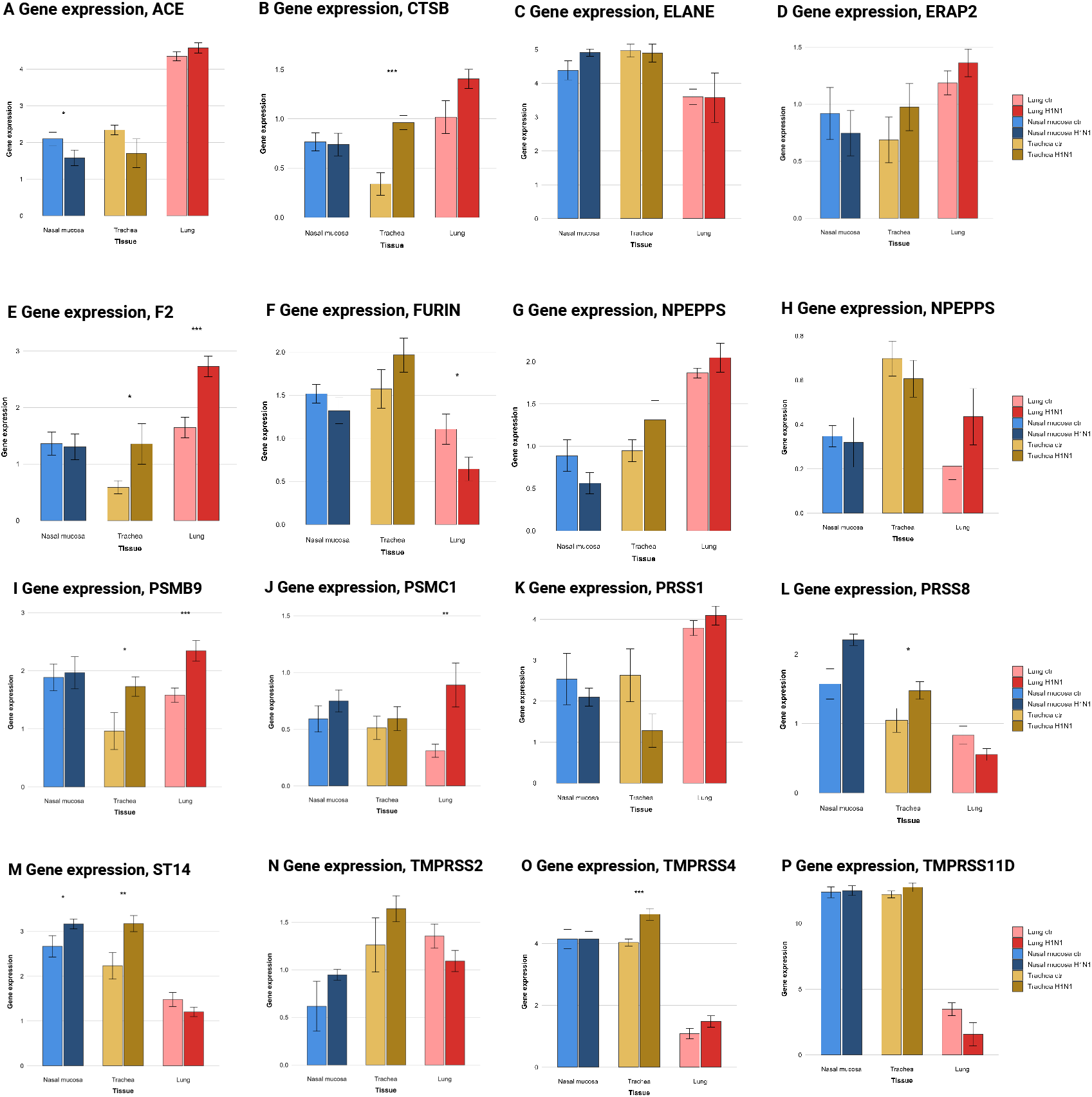
Protease expression, microfluidic qPCR

## References

1. Carter, T. & Iqbal, M. The influenza a virus replication cycle: A comprehensive review, DOI: 10.3390/v16020316 (2024).

2. Mair, C. M., Ludwig, K., Herrmann, A. & Sieben, C. Receptor binding and ph stability — how influenza a virus hemagglutinin affects host-specific virus infection. Biochimica et Biophys. Acta (BBA) - Biomembr. 1838, 1153–1168, DOI: 10.1016/J.BBAMEM.2013.10.004 (2014).

3. Pampalakis, G., Zingkou, E., Panagiotidis, C. & Sotiropoulou, G. Kallikreins emerge as new regulators of viral infections. Cell. Mol. Life Sci. 78, 6735–6744, DOI: 10.1007/s00018-021-03922-7 (2021).

4. Leu, C.-H. et al. Kallistatin ameliorates influenza virus pathogenesis by inhibition of kallikrein-related peptidase 1-mediated cleavage of viral hemagglutinin. Antimicrob. Agents Chemother. 59, 5619–5630, DOI: 10.1128/AAC.00065-15 (2015).

5. Peitsch, C., Klenk, H.-D., Garten, W. & Böttcher-Friebertshäuser, E. Activation of influenza a viruses by host proteases from swine airway epithelium. J. Virol. 88, 282–291, DOI: 10.1128/JVI.01635-13 (2014).

6. Böttcher-Friebertshäuser, E. et al. Cleavage of influenza virus hemagglutinin by airway proteases tmprss2 and hat differs in subcellular localization and susceptibility to protease inhibitors. J. Virol. 84, 5605–5614, DOI: 10.1128/jvi.00140-10 (2010).

7. Böttcher, E. et al. Proteolytic activation of influenza viruses by serine proteases tmprss2 and hat from human airway epithelium. J. Virol. 80, 9896–9898, DOI: 10.1128/jvi.01118-06 (2006).

8. Limburg, H. et al. Tmprss2 is the major activating protease of influenza a virus in primary human airway cells and influenza b virus in human type ii pneumocytes. J. Virol. 93, DOI: 10.1128/jvi.00649-19 (2019).

9. Böttcher-Friebertshäuser, E., Garten, W. & Klenk, H. D. Activation of viruses by host proteases (Springer International Publishing, 2018).

10. Lubinski, B. & Whittaker, G. R. Host cell proteases involved in human respiratory viral infections and their inhibitors: A review, DOI: 10.3390/v16060984 (2024).

11. Xia, Q., Liu, X. & Huang, H. Host proteases: key regulators in viral infection and therapeutic targeting. Front. Immunol. 16, DOI: 10.3389/fimmu.2025.1671173 (2025).

12. Laporte, M. & Naesens, L. Airway proteases: an emerging drug target for influenza and other respiratory virus infections, DOI: 10.1016/j.coviro.2017.03.018 (2017).

13. Sadler, A. J. & Williams, B. R. G. Interferon-inducible antiviral effectors. Nat. Rev. Immunol. 8, 559–568, DOI: 10.1038/nri2314 (2008).

14. Zhang, M. et al. Isgylation in innate antiviral immunity and pathogen defense responses: A review. Front. Cell Dev. Biol. 9, DOI: 10.3389/fcell.2021.788410 (2021).

15. Oshiumi, H. et al. Ddx60 is involved in rig-i-dependent and independent antiviral responses, and its function is attenuated by virus-induced egfr activation. Cell Reports 11, 1193–1207, DOI: 10.1016/j.celrep.2015.04.047 (2015).

16. Rawlings, N. D. et al. The merops database of proteolytic enzymes, their substrates and inhibitors in 2017 and a comparison with peptidases in the panther database. Nucleic Acids Res. 46, D624–D632, DOI: 10.1093/nar/gkx1134 (2018).

17. Lange, P. F. & Overall, C. M. Topfind, a knowledgebase linking protein termini with function. Nat. Methods 8, 703–704, DOI: 10.1038/nmeth.1669 (2011).

18. Ahmad, S. et al. The uniprot website api: facilitating programmatic access to protein knowledge. Nucleic Acids Res. 53, W547–W553, DOI: 10.1093/nar/gkaf394 (2025).

19. Bond, J. S. Proteases: History, discovery, and roles in health and disease. J. Biol. Chem. 294, 1643–1651, DOI: 10.1074/JBC.TM118.004156 (2019).

20. Wong, J., Layton, D., Wheatley, A. K. & Kent, S. J. Improving immunological insights into the ferret model of human viral infectious disease. Influ. Other Respir. Viruses 13, 535–546, DOI: 10.1111/irv.12687 (2019).

21. Pathak, T., Pal, S. & Banerjee, I. Cathepsins in cellular entry of human pathogenic viruses. J. Virol. 99, DOI: 10.1128/jvi.01642-24 (2025).

22. Bertram, S. et al. Tmprss2 and tmprss4 facilitate trypsin-independent spread of influenza virus in caco-2 cells. J. Virol. 84, 10016–10025, DOI: 10.1128/jvi.00239-10 (2010).

23. Tanaka, Y. L. et al. Generation of a porcine cell line stably expressing pig tmprss2 for efficient isolation of swine influenza virus. Pathogens 13, 18, DOI: 10.3390/pathogens13010018 (2023).

24. Zanella, G. C. et al. Pigs lacking tmprss2 displayed fewer lung lesions and reduced inflammatory response when infected with influenza a virus. Front. Genome Ed. 5, DOI: 10.3389/fgeed.2023.1320180 (2024).

25. Kwon, T. et al. Gene editing of pigs to control influenza a virus infections. Emerg. Microbes & Infect. 13, DOI: 10.1080/22221751.2024.2387449 (2024).

26. Kristensen, C. et al. Experimental infection of pigs and ferrets with “pre-pandemic,” human-adapted, and swineadapted variants of the h1n1pdm09 influenza a virus reveals significant differences in viral dynamics and pathological manifestations. PLoS Pathog. 19, DOI: 10.1371/journal.ppat.1011838 (2023).

27. Laybourn, H. A. et al. Tracking mucosal innate immune responses to three influenza a virus strains in a highly translational pig model using nasopharyngeal swabs. Innate Immun. 31, DOI: 10.1177/17534259251331385 (2025).

28. Yadati, T., Houben, T., Bitorina, A. & Shiri-Sverdlov, R. The ins and outs of cathepsins: Physiological function and role in disease management. Cells 9, 1679, DOI: 10.3390/cells9071679 (2020).

29. Ruiz-Blázquez, P. et al. Cathepsin d is essential for the degradomic shift of macrophages required to resolve liver fibrosis. Mol. Metab. 87, 101989, DOI: 10.1016/j.molmet.2024.101989 (2024).

30. Solberg, R. et al. The mammalian cysteine protease legumain in health and disease. Int. J. Mol. Sci. 23, 15983, DOI: 10.3390/ijms232415983 (2022).

31. Dinesh, R. K. et al. Membrane-wide screening identifies potential tissue-specific determinants of sars-cov-2 tropism. PLOS Pathog. 21, e1013157, DOI: 10.1371/journal.ppat.1013157 (2025).

32. Mena, I. et al. Origins of the 2009 h1n1 influenza pandemic in swine in mexico. eLife 5, DOI: 10.7554/eLife.16777 (2016).

33. Weng, S. S. et al. Sensitive determination of proteolytic proteoforms in limited microscale proteome samples. Mol. Cell. Proteomics 18, 2335–2347, DOI: 10.1074/mcp.TIR119.001560 (2019).

34. Kalogeropoulos, K. et al. Clipper 2.0: Peptide level annotation and data analysis for positional proteomics. DOI: 10.1101/2023.11.30.569335.

35. Thomsen, M. C. F. & Nielsen, M. Seq2logo (2012).

36. Perez-Riverol, Y. et al. The PRIDE database at 20 years: 2025 update. Nucleic acids research 53, D543–D553, DOI: 10.1093/nar/gkae1011 (2025).

